# Intracellular anti-leishmanial effect of Spergulin-A, a triterpenoid saponin of *Glinus oppositifolius*

**DOI:** 10.1101/458653

**Authors:** Saswati Banerjee, Niladri Mukherjee, Rahul L. Gajbhiye, Parasuraman Jaisankar, Sriparna Datta, Krishna Das Saha

**Affiliations:** Cancer Biology and Inflammatory Disorder Division, CSIR-Indian Institute of Chemical Biology, 4 Raja S. C. Mullick Road, Kolkata-700032, India; Organic and Medicinal Chemistry Division, CSIR-Indian Institute of Chemical Biology, 4, Raja S.C. Mullick Road, Kolkata 700032, India; Department of Chemical Technology, University of Calcutta, Kolkata 700009, India

**Keywords:** Visceral leishmaniasis, *Glinus oppositifolius*, Spergulin-A, Macrophage, Immunostimulation, Anti-amastigote

## Abstract

Many of present chemotherapeutics are inadequate and also resistant against visceral leishmaniasis (VL) an immunosuppressive ailment caused by *Leishmania donovani*. Despite the interest in plant-based drug development, very few number of antileishmanial drugs from plant source is available. *Glinus oppositifolius* had been reported in favor of being immunomodulators along with other traditional uses. Novel anti-VL therapies can rely on host immune-modulation with associated leishmanicidal action. With this rationale, an n-BuOH fraction of the methanolic extract of the plant and obtained triterpenoid saponin Spergulin-A were evaluated against acellular and intracellular *L. donovani*. Immunostimulatory activity of them was confirmed by elevated TNF-α and extracellular NO production from treated MФs and was found nontoxic to the host cells. Identification and structure confirmation for isolated Spergulin-A was performed by ESI-MS, ^13^C, and ^1^H NMR. Spergulin-A was found ineffective against the acellular forms while, against the intracellular parasites at 30μg/ml, the reduction was 92.6% after 72h. Spergulin-A enhanced ROS and nitric oxide (NO) release and changes in Gp91-phox, i-NOS, and pro and anti-inflammatory cytokines elaborated its intracellular anti-leishmanial activity. The results supported that *G. oppositifolius* and Spergulin-A can potentiate new lead molecules for the development of alternative drugs against VL.

## Introduction

Visceral Leishmaniasis (VL) considered as the most severe form of leishmaniasis caused by *Leishmania donovani* and untreated patients nearly always die [1]. Leishmaniasis is endemic in nearly 100 countries with approximately 350 million people are at risk with an estimated yearly incidence of 500,000 cases and almost 70,000 deaths. Leishmaniasis is liable for the ninth most substantial infectious diseases burden, however, is mostly disregarded among tropical disease priorities [2]. Sodium antimony gluconate, Miltefosine, Pentamidine, and Amphotericin B are the primary therapeutics though associated with toxicity or resistance [3].

Pathogen recognition by neutrophils, macrophages, dendritic cells, and natural killer cells activates intracellular signaling pathways leading into inflammatory responses like inflammasome activation and IL-1β production, which is not the case for leishmanial infection [4]. *L. donovani* infection is characterized by the parasite-induced active subversion of the host immune system and also immune deviation which additively favors infection establishment [5]. Additionally, *Leishmania* prevents inflammatory response by the impaired release of different pro-inflammatory cytokines (IL-1, TNF-α, IL-12) and enhanced releases of immunosuppressive signaling molecules, such as arachidonic acid metabolites and the cytokines TGF-β and IL-10 [6]. Involvement of co-stimulatory molecule B7-CTLA-4 is associated with increased TGF-ß production in VL with increased apoptosis of CD4^+^ T cells and decreased macrophage apoptosis [7]. Leishmanial infection also undermines a generation of microbicidal macrophage nitric oxide (NO) and reactive oxygen species (ROS) with the hindrance of antigenic peptide display to T cells, and permeation of IL-10 producing T regulatory cells [8]. In recent years increased instances of VL has been reported in connection with immune-suppressed AIDS patients [8].

Discovery of novel compounds which intercedes host immune-modulation associated with leishmanicidal function having permissible side effects is a precise research objective [9,10]. It is also of vital importance that the drug required for parasite elimination in immune-stimulated cells was significantly less than the immune-suppressed ones [11].

The anti-parasitic effectiveness of plant extracts majorly relies upon secondary metabolites of diverse chemical groups including alkaloids, polyphenol, flavonoids, terpenoids, phenylpropanoids, etc. [12]. For isolation and characterization of a herbal extract or an active compound, different research strategies can be employed mainly in the extraction steps. However, bioactivity-guided fractionation considered as simple, rapid, cost-effective and reproducible [1]. For obtaining a potent anti-leishmanial agent with immune-modulation, different studies were conducted with compounds like aslicarin A, niranthrin, skimmianine, quassin, tannins, linalool, etc. nevertheless with varying degree of effectiveness and satisfaction [8].

Aerial parts of *Glinus oppositifolius* (Family Molluginaceae) are used for treating abdominal pain and jaundice, while decoction is used against malaria [13]. Plants of this genus were previously documented for the presence of triterpenoid saponins [14], and isolated pectin polysaccharides are antiprotozoal [15] and immunomodulators [14]. *G. oppositifolius* is indicated for wound healing and used in traditional medicine for treating diarrhea, joint pains, inflammations, intestinal parasites, fever, boils and skin disorders [16]. Some triterpenoid saponins, 3-O-(β-D-xylopyranosyl)-spergulagenin-A, Spergulacin, Spergulacin-A Spergulin-A, and Spergulin-B had been isolated from *G. oppositifolius* [17]. Application of plant-based immunomodulators is smart as they mediate their effectiveness by enhancing the inherent host-derived protective machinery without the involvement of specific microbicidal agents namely antibiotics [13].

Most of the available antimalarial drugs are plant-derived; regrettably, there is no anti-leishmanial drug present which is of plant origin. The recent efforts to achieve this goal also were restricted against the promastigotes [18]. In the present work, an attempt has been taken to evaluate the intracellular anti-leishmanial activity of this plant and its bioactive component, Spergulin-A emphasizing on the immunostimulatory activity.

## Materials and methods

### Chemicals and Reagents

Cell culture media, serum, antibiotics, HEPES (Gibco, USA), CFSE, DAPI (Invitrogen, USA), MTT, Miltefosine, DMSO (Sigma Chemical Co. USA), cytokine Assay kit (Thermo Scientific, USA), DAF-2 DA, nitric oxide assay kit, DCFDA (Calbiochem, USA) and all other chemicals were of the highest grade commercially available. Primary and secondary antibodies were obtained from Santa Cruz Biotechnology (USA) or Cell signaling technologies (USA).

### Isolation of methanolic fractions from *G. oppositifolius*

The aerial parts of *G. oppositifolius* were shade dried (1kg) and were first defatted using petroleum ether (60-80°C, 3.5L X 3) at room temperature for 48h. The marc was than subjected to extraction using MeOH (3.5L X 3) at room temperature for 48h. The extract was then filtered and MeOH was evaporated under reduced pressure and finally was lyophilized to obtain the crude MeOH extract (13g). A part (10g) of ME was then suspended in milli-Q water and partitioned sequentially with EtOAc and n-BuOH. Each fraction was evaporated under vacuum and lyophilized to yield the EtOAc fraction (EAF; 3.6g), n-BuOH fraction (NBF; 4.1g) and aqueous fraction (AF; 2.3g). All the fractions were stored at 4ºC till further use.

Among these four fractions, n-BuOH showed promising antileishmanial activity. Around 4g of NBF was subjected to column chromatography of Diaion HP 20 (100g) and the column washed with water followed by 30, 40, 60, 80 and 100% of MeOH to obtain a total of six fractions. Fractions eluted with 50% MeOH showed similar spots on TLC and were mixed and then re-chromatographed on Dianion HP 20 column to furnish 7mg of Spergulin-A. The compound isolation procedures were performed as followed by Kumar et al. [19].

### Isolation and characterization of Spergulin-A

ESI Mass spectra were recorded on an Agilent 6545 Q-TOF mass spectrometer System. ^1^H and ^13^C NMR were recorded on a Bruker Ultrashield NMR (600 MHz) in pyridine-d5 with TMS as an internal standard. Diaion HP 20 was used for column chromatography; silica gel (60 F254) was used for TLC and spots were visualized by spraying with Lieberman–Burchard reagent followed by heating.

### Parasite culture and maintenance

Passage in BALB/c mice maintained *Leishmania donovani* [MHOM/IN/1983/AG83] infection. Complete M199 media (10% FBS, pH 7.4, 100U/ml penicillin G-sodium, 100μg/ml streptomycin sulfate, 25mM HEPES) was used to culture the promastigotes. Axenic amastigote from the log phase culture of promastigotes was prepared as described by Saar et al. [20].

### Macrophage culture, parasite infection, and treatment

RAW 264.7 MФ cell line was obtained from the ATCC and maintained in complete RPMI 1640 (10% FBS, 100μg/ml streptomycin sulfate, 100U/ml penicillin G sodium, 0.2% sodium bicarbonate, 25 mM HEPES) in a humidified atmosphere and 5% CO_2_ at 37°C.

Infection of MФs with *L. donovani* promastigotes was performed in a ratio of 1:10 (MΦ: parasite) for 4h then washed twice with media. Normal and parasitized macrophages were treated with *G. oppositifolius* fractions or Spergulin-A at required concentrations up to 72h. Infection was measured by counting the intracellular parasites and expressed as parasite count/20 MΦs. Miltefosine served as positive anti-leishmanial reference.

### ELISA assay

Cell-free supernatants from MФs (1×10^5^ cells/well) were collected, and the concentration of TNF-α was estimated by sandwich ELISA, using a commercially available assay kit [21].

### MTT assay

20μl of MTT (5mg/ml in PBS) was added to each well of the control and treated MФs (4×10^3^) and 5×10^3^ parasites (promastigote and axenic amastigote) in a 96 well plate and incubated for 4h at 37°C and the formazan crystals were dissolved in 150μl of DMSO. Absorption was measured at 595nm by an ELISA reader [22].

### Measurement of extracellular NO

The collected supernatants from control, infected and treated MФs were incubated with equal volumes of Griess reagent, NED (0.1% in distilled water) and Sulphanilamide (1% in 5% H_3_PO_4_) at room temperature for 10min. The absorbance was measured at 540nm on a microplate reader. NO concentration was determined using a dilution of sodium nitrite as the standard [21].

### FACS and Confocal microscopy with CFSE tagged L. donovani

MФs were parasitized with CFSE-tagged (25μM/1×10^6^ promastigote in 1ml media for 30min) promastigotes followed by Spergulin-A (10,20,30μg/ml) treatment for 24h.

For confocal microscopy, MФs (1×10^5^/glass coverslips) after infection and treatment, washed twice with PBS, fixed in chilled 70% ethanol and nuclei were stained with DAPI and observed under Olympus Fluoview FV10i confocal microscope with 60X objective lens. For FACS, MФs (1×10^5^/well in a six-well plate) after infection and treatment, washed twice with PBS then harvested by scraping and re-suspended in 400μl PBS and analyzed by BD FACS LSR Fortessa with excitation at 494nm and emission at 518nm.

### FACS analyses of intracellular ROS and NO

The level of intracellular ROS was determined based on the change in fluorescence of H2DCFDA. DAF-2 DA was used for detecting intracellular NO [3]. Briefly, after infection and treatment, MФs were scrapped and incubated in PBS containing DAF-2DA (7.0μM) at 37°C for 30min or 20μM DCF-DA at 37°C for 15min and then analyzed by BD FACS LSR Fortessa with excitation at 480nm and emission at 515nm for both.

### Western blot analysis

40µg of proteins harvested from MФs were electrophoretically separated in SDS-polyacrylamide gel and transferred to PVDF membrane, blocked with BSA and incubated with respective primary antibodies overnight. The membranes were then incubated with HRP conjugated secondary antibodies, and immunoreactive bands were visualized by adding proper substrates. β-actin was used as loading controls [23].

### Statistical analysis

All values were expressed as mean ± SEM obtained from at least three replicate experiments. Statistical significance and differences among groups were assessed with One-Way analysis of variance (ANOVA) followed by Dunnett’s test. P values ≤ 0.05 (*) or ≤ 0.01 (**) were considered as indicative of significance.

## Results

### Immunostimulatory effect of methanolic extract of *G. oppositifolius*

Of the aqueous, ethyl acetate and n-BuOH (50:50) fractions of *G. oppositifolius* methanolic extract, only the n-BuOH fraction (50μg/ml) showed a considerable increase in TNF-α and extracellular NO production (Supple. Figure 1) from the treated (24h) MФs. The six sub-fractions (50μg/ml) of the n-BuOH fraction were also checked for the same parameters (Figure 1A and B), and highest augmentation was noticed in fraction 4 for being a worthy immunostimulatory agent and the lead fraction.

**Fig. 1.**
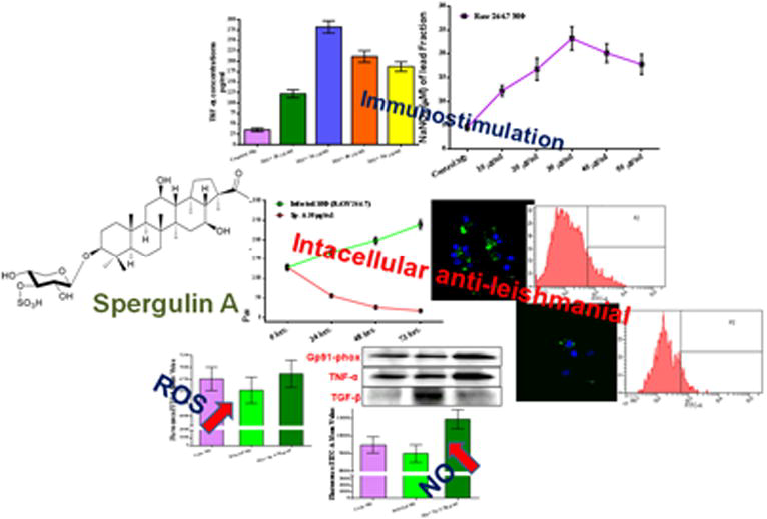
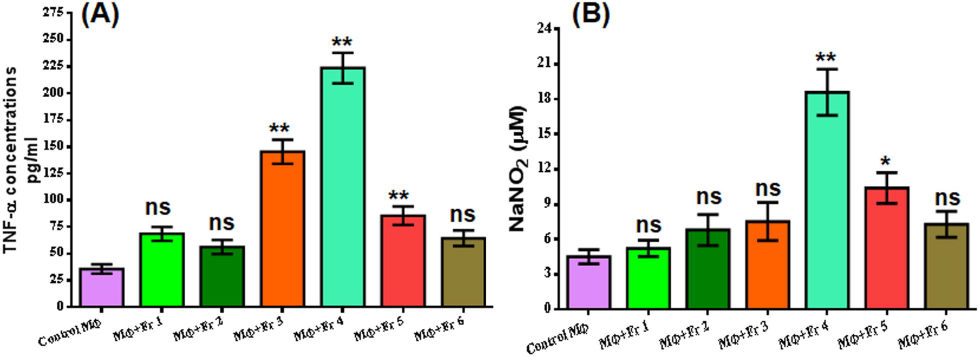
Evaluation of immunostimulation by subfractions of n-BuOH fraction of *G. oppositifolius* MeOH extract. (A) The subfractions (50μg/ml) were checked for altered TNF-α production after 24h treatment of RAW 264.7 MФs from the culture supernatant. (B) Extracellular NO production was monitored in the culture supernatant of treated (six subfractions) and control after 24h by Griess reagent. All values are expressed as mean ± SEM from triplicate assays from three independent experiments (P values ≤ 0.05 (*) or ≤ 0.01 (**) vs. control).

### Dose-dependent immunostimulatory effect of the lead fraction

A dose-dependent increase in TNF-α production was monitored in treated (24h) MФs in two sets (Figure 2B and C). In first set the lead fraction was applied at the doses of 10, 20, 40 and 80μg/ml (Figure 2B) and in the second set at the doses of 20, 30, 40 and 50μg/ml and most increments was noticed at 30μg/ml (Figure 2C) and selected as the significant dose for this study.

**Fig. 2.**
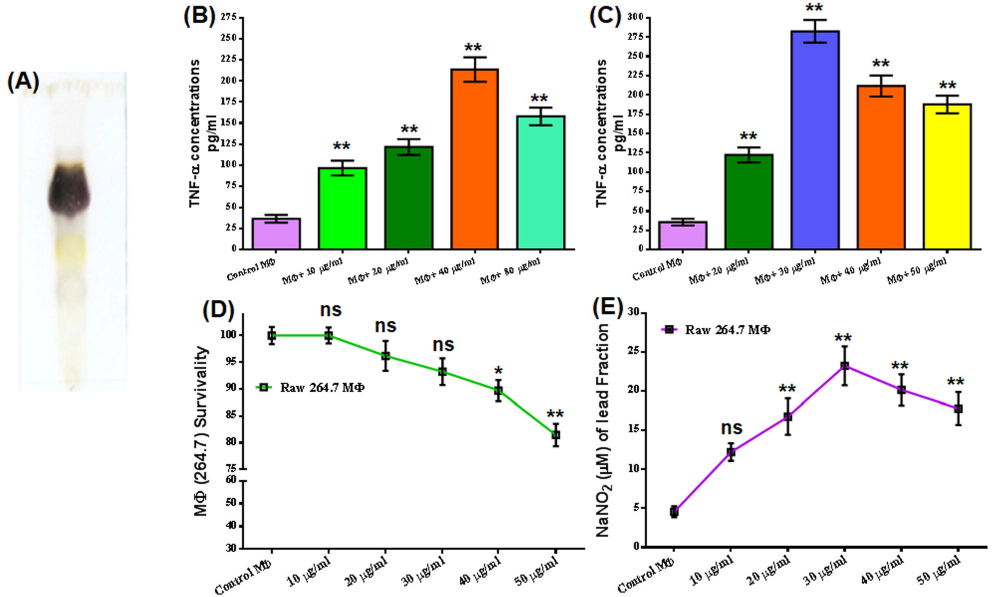
Dose-dependent evaluation of the immunostimulatory effect of the n-BuOH lead fraction and impact on RAW 264.7 MФ survival. (A) TLC profile of n-BuOH lead fraction. (B, C) The dose-dependent release of TNF-α was monitored in treated (24h) RAW 264.7 MФs, and highest increment was found in 30μg/ml. (D) Survival of RAW 264.7 MФs at different doses of an n-BuOH lead fraction after 24h exposure measured by MTT assay. (E) Dose-dependent evaluation of extracellular NO production was monitored in the culture supernatant of 24h treated RAW 264.7 MФs by Griess reagent. All values are denoted as mean ± SEM from triplicate assays from three independent experiments (P values ≤ 0.05 (*) or ≤ 0.01 (**) vs. control).

### Assessment of macrophage survival and extracellular NO release

Survival of the MФs after exposing them to increasing concentrations (10,20,30,40 and 50μg/ml) of the lead fraction for 24h was monitored by MTT assay (Figure 2D). The lead fraction was found safe for this dose range, and survival was about 81.45% even at 50μg/ml. Most increment (5fold) in extracellular NO was noticed for 30μg/ml of the lead fraction (Figure 2E). The increase was also significant for most of the doses.

### Identification and characterization of Spergulin-A

Among those six n-BuOH bioactive sub-fractions, fraction 4 have shown most significant upliftment in TNF-α and extracellular NO production. After purification of this fraction by using column chromatography, a white amorphous compound was obtained. Molecular formula C_35_H_58_O_11_S (Figure 3A) was assigned from the ESI-MS (Supple. Figure 2). Structural elucidation of this compound was achieved by critical analysis of the ^1^H NMR, ^13^C NMR and ^13^C NMR DEPT results (Supple. results and Supple. Figure 3–6). All the spectral data of this compound were found to be in complete agreement with reported ones and structure was then confirmed as Spergulin-A [24].

**Fig. 3.**
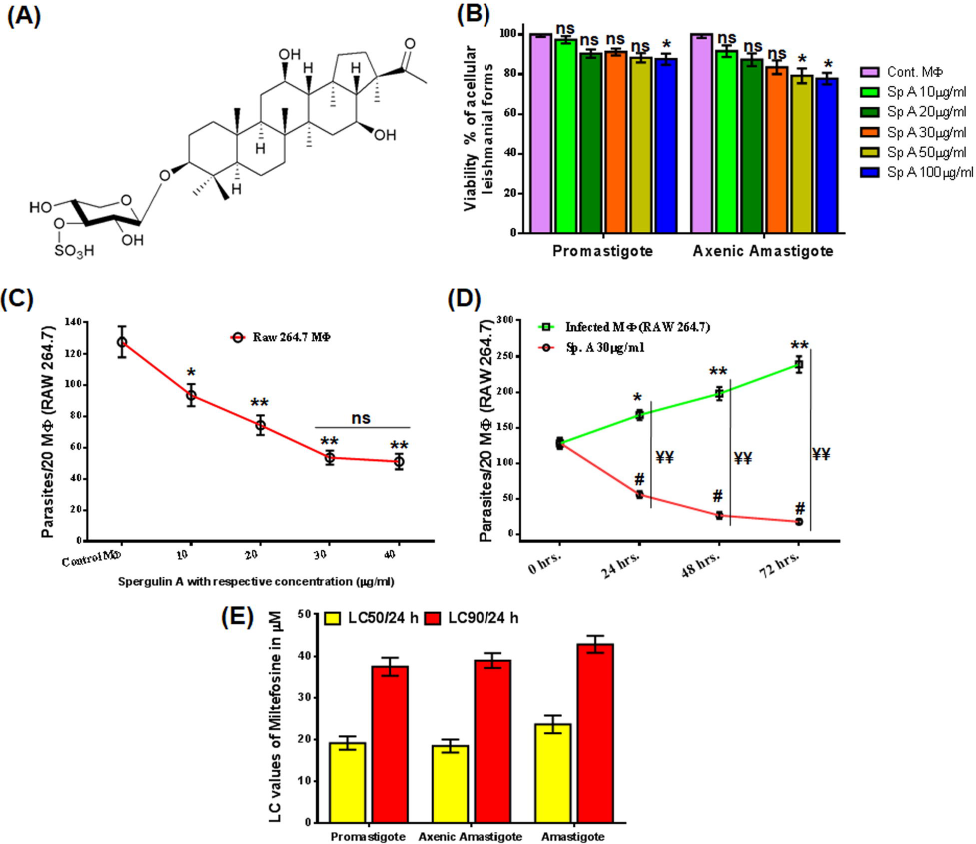
Anti-leishmanial activity of Spergulin-A. (A) Structure of the triterpenoid saponins Spergulin-A. (B) Dose-dependent effects of Spergulin-A against acellular forms namely the promastigotes and axenic amastigote of *L. donovani* after 24h. (C) Dose-dependent effect of Spergulin-A against the intracellular amastigote of *L. donovani* after 24h of treatment and expressed as number of parasites/20 MФs. (D) 24h time lapse evaluation of anti amastigote effect of Spergulin-A at 30μg/ml for 72h. (E) Miltefosine LC_50_/24h and LC_90_/24h values against promastigote, axenic amastigote and intracellular amastigote forms of *L. donovani*. All values are expressed as mean ± SEM from triplicate assays from three independent experiments (P values ≤ 0.05 (*) or ≤ 0.01 (**) vs. control).

### Assessment of anti-leishmanial activity of Spergulin-A

#### Effect on different forms of *L. donovani*

Anti-leishmanial effect of Spergulin-A (10,20,30,50,100μg/ml) was first evaluated by MTT assay against promastigote and axenic amastigote, the acellular forms (Figure 3B) after 24h. Viability was 87.6% and 77.8% respectively for promastigote and axenic forms even at 100μg/ml. At the lower doses (10, 20 and 30μg/ml) the viability reduction was found to be non-significant (Figure 3B).

After that, the effect of Spergulin-A was evaluated against the MФ internalized parasites (Figure 3C). Here, a significant reduction of the parasite was noticed as low as 10μg/ml and further increased with the increment of doses (Figure 3C). However, above 30μg/ml MФ internalized antileishmanial effect of Spergulin-A had reached a plateau, and thus further experiments were mostly conducted with 30μg/ml.

#### Time-dependent intracellular leishmanicidal effect

Efficacy of Spergulin-A (30μg/ml) against the intracellular parasites was evaluated at every 24h interval up to 72h (Figure 3D). Inside the infected MФs parasite count increased significantly. The treated MФs exhibited significantly reduced numbers of parasites from the initial infection as well as the corresponding observation point of infected macrophages (Figure 3D). The parasite reduction at 72h was 86.2% less than the initial point of infection and 92.6% less than the 72h count of infected MФs. Miltefosine was used in parallel as a positive reference anti-leishmanial compound and LC_50_/24h and LC_90_/24h values against these three forms were expressed as histogram (Figure 3E).

#### Dose-dependent quantitative and qualitative anti-leishmanial effect

After infecting the MФs with CFSE-tagged promastigotes and treatment with Spergulin-A (10, 20 and 30μg/ml, 24h), the cells were evaluated by FACS in FITC filter (Figure 4A). Highest intensity was noticed in the infected panels (Figure 4A.ii) and with the increment of Spergulin-A doses the intensity gradually reduces which is reciprocal to the reduced parasite count (Figure 4A.iii-v). Interestingly, no significant changes in number of parasitized MФs observed within the treatment panel (Figure 4A.vi) and what changes were the intensity denoted the parasite count (Figure 4A.vii). Thus, possibly Spergulin-A reduced the parasite by immune-stimulating the host MФs.

**Fig. 4.**
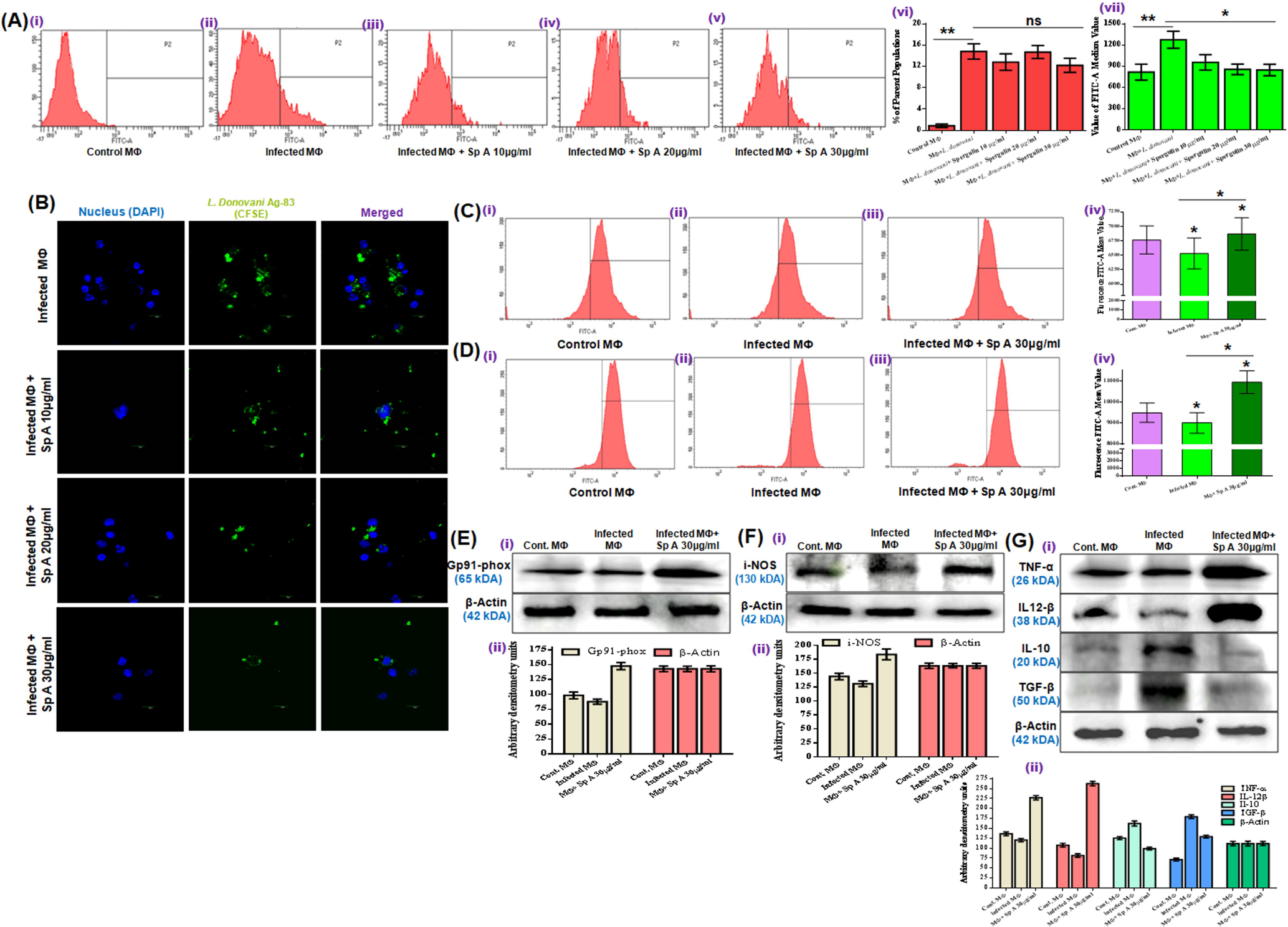
Elaboration of the Anti-leishmanial activity of Spergulin-A. (A) Quantitative dose-dependent assessment of intracellular CFSE stained antileishmanial activity of Spergulin-A by flow cytometry after 24h treatment. (B) Confocal laser-scanning micrographs to evaluate the reduction in intracellular CFSE stained *L. donovani* after 24h of Spergulin-A treatment (magnification 120X). (C) FACS monitored the changes in ROS production in control, infected, and treated (30μg/ml Spergulin-A) MФs with H2DCFDA. (D) The changes in intracellular NO production were monitored by FACS with fluorescent probe DAF-2 DA. (E, F, G) Changes in the expression of Gp91-Phox, i-NOS and cytokines namely pro-inflammatory TNF-α, IL-12β and anti-inflammatory IL-10 and TGF-β with β-actin as loading control and densitometry analyses was provided after band normalization. Images are representative of three separate experiments, and all values are expressed as mean ± SEM from triplicate assays from three independent experiments (P values ≤ 0.05 (*) or ≤ 0.01 (**) vs. control).

Confocal micrographs depicted the reduction in parasite emitting green fluorescence with the increment of Spergulin-A doses. Moreover, most decreases were noticed when the parasitized MФs was treated with 30μg/ml of it (Figure 4B).

#### Intracellular ROS and NO status due to infection and treatment

The fluorescence intensity for H2DCFDA reciprocal to ROS production was found to be minimal for the infected panel (Figure 4C.ii) and significantly higher in the treated panel (Figure 4C.iii). The fluorescence intensity of DAF-2DA for NO was also found to be highest for the treated panel (Figure 4D.iii) compared to the control and infected MФs (Figure 4D.i and ii).

#### Estimation of alteration of different cytokines and immunostimulators

In western blot analysis, it was observed that Gp91-phox liable for anti-microbial ROS production in MФs got up regulated in treated panel (Figure 4E). Level of i-NOS which regulates anti-leishmanial NO production also increased with Spergulin-A treatment compared to control and infected MФs (Figure 4F).

Expression of pro-inflammatory TNF-α and IL-12β got down-regulated with infection and considerably up-regulated in treatment panel (Figure 4G). Anti-inflammatory IL-10 and TGF-β expression were up-regulated in infected MФs compared to the control, and with the treatment of Spergulin-A (30μg/ml) their appearance got normalized like that of control (TGF-β) or even less (IL-10) (Figure 4G).

## Discussion

Precisely an immune-suppressive ailment, VL inhabit and modulate the microbicidal function of the macrophages and create a microenvironment favoring parasite growth inside the visceral organs by modulating pro and anti-inflammatory cytokines and impairing ROS and NO release [6]. The present *in vitro* study aimed to assess the immunostimulatory property of *G. oppositifolius* and isolation of a compound Spergulin-A which can reduce the intracellular parasites by immune-stimulation within a safe dose range for the host cells. Primary appraisal of an extract that can have an immunostimulatory effect was based upon the increased release of TNF-α and extracellular NO from treated MФs. It was observed that at an applied same dose (50μg/ml, 24h) only the n-BuOH fraction showed a considerable increase in TNF-α and extracellular NO production. The selection of TNF-α and extracellular NO as the potent immunostimulatory index in the present context is vital because endogenous TNF-α produced by infected macrophages can elicit the release of L-arginine-derived nitrogen intermediates detrimental for the intracellular parasites [9]. The six n-BuOH sub-fractions were again verified for immune-stimulation, and fraction 4 emerged as the most promising one. The fraction was also considered to be safe for the MФs even at 50μg/ml (81.5% viability) having CC_50_ well above 100μg/ml.

Interested by the initial observations on elevated immunostimulation by the n-BuOH sub-fraction 4 and based upon the previous inspection of the presence of triterpenoid saponins in this fraction [24] a triterpenoid saponin Spergulin-A was isolated and purified following the well-established method (Fig. 3A and Supple.information) [24,19].

Evaluation of the anti-leishmanial property of Spergulin-A was the principal research interest after identification and isolation of it. The possible anti-leishmanial effect was first checked against the acellular forms of the parasite but without major success even at 100μg/ml (Figure 3B). Therefore, if Spergulin-A has any leishmanicidal effect on the intracellular parasites that will possibly mediate by altering the microenvironment of the host MФs. After 24h of incubation Spergulin-A (30μg/ml) was found to be pretty useful in reducing the intracellular parasite count and when applied for a prolonged period of 72h reduction of intracellular *Leishmania* was significant in respect to both the infected MФs of corresponding time point and initial parasitized MФs (Figure 3C,D). It is also of great significance that no considerable alteration in number of parasitized MФs was found in treatment panel while reduction of CFSE-stained internalized parasites was notable (Figure 4A). This specific observation strongly advocated in favor of immunostimulation by Spergulin-A which mediated parasite killing. For being leishmanicidal, Spergulin-A should be useful in enhancing the release of NO and ROI, well recognized for their worth against *Leishmania* [6,25]. NO is especially critical for intracellular parasite clearance as it was reported that mice with impaired inducible nitric oxide synthase (i-NOS) and thus restricted production and release of NO are incapable of *Leishmania* control even for isolated macrophages *in vitro* [26]. At the dose of 30μg/ml, Spergulin-A, showed enhanced production of intracellular NO and ROI (Fig. 4C,D) which was found to be directly reciprocal to the reduction of intracellular parasites and signified their connection. In the western blot analysis also increment in i-NOS production (Figure 4F) was noticed in Spergulin-Atreated parasitized MФs in amendable enhanced NO production and parasite control. The same interpretation is also pertinent for an increase of Gp91-phox, a subunit of the NADPH oxidase and enhancement of ROI (Figure 4 E,C) in Spergulin-A mediated intracellular parasite killing. It was previously established that intracellular leishmanial killing proceeds by NO production from arginine by i-NOS, and superoxide (O2^−^) generated by the NADPH oxidase [27]. Several Leishmania species induce immunosuppressive TGF-β production, and diminution of TGF-β secretion is in direct correlation with enhanced i-NOS production [6] and can lead to internalized parasite removal as found with Spergulin-A treatment. TGF-β also found to increase the VL progression and averts disease cure in murine models [27]. IL-10 is another anti-inflammatory cytokine which makes macrophages indifferent to various activation signals and thus causes impairment of intracellular parasite killing also by down-regulating the production of TNF-α and NO [28]. IL-10 production increased in infected macrophages *in vitro*, apparently via interaction with the Fc receptor [29] and which down-regulated with Spergulin-A treatment (Figure 4G) in connection with parasite reduction. Parasite infection is responsible for the suppression of macrophage microbicidal activity that relies upon NO, ROI and cytokines like IL-1, IL-12β, and TNF-α [30] as with infection the levels of TNF-α IL-12β got down-regulated, but with the application of Spergulin-A, these cytokines significantly elevated which signifies their role in intracellular *Leishmania* parasite control.

## Conclusion

The inconsistency of direct leishmanicidal effect of *G. oppositifolius* and Spergulin-A isolated from it against promastigotes and axenic amastigote on the one hand and the proved efficacy of them against intracellular parasites was explained by emphasizing host MФ immunostimulation as elaborated precisely in the present study. Bio-guided fractionation was employed to isolate and identify Spergulin-A as the immunostimulant, and its anti-leishmanial property was evaluated at a dose best suited for the research and found to be safe against the host cells assessed *in vitro* in RAW 264.7 MФs infected with a virulent strain of *L. donovani*.

## Acknowledgment

Sincere thanks are given to Science and Engineering Research Board, Govt. of India (Grant No. PDF/2016/001437 and EMR/2015/001674) for financial assistance. The manuscript has been checked critically by the full version of the Grammarly software and corrected by Dr. Basudeb Achari, ex-scientist of CSIR-Indian Institute of Chemical Biology.

