## Supplementary File for "Intracellular anti-leishmanial effect of Spergulin-A, a triterpenoid saponin of *Glinus oppositifolius*"

**Content:**

NMR data of Spergulin A

Supplementary Figures: Fig. 1-6

Reference

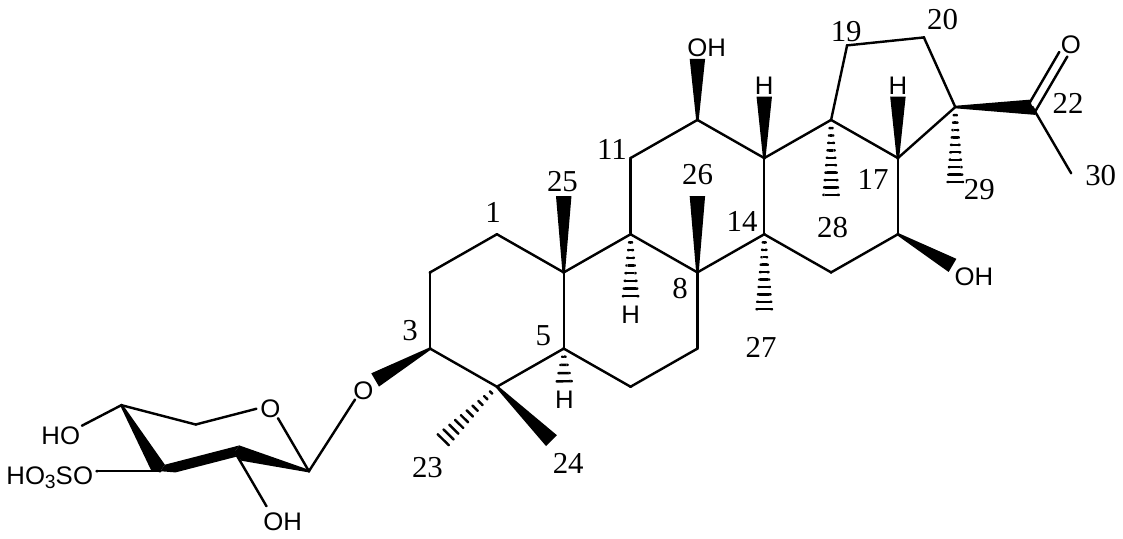

**Structure of Spergulin A**

**NMR data of Spergulin A**

White amorphous solid; ESI-MS *m/z*: 731.34 [M+2Na-H]^+^, (calc. for C_35_H_57_O_11_S, *m/z* 685.36216). ^1^H NMR (C_5_D_5_N; 600 MHz). 0.61, 0.78, 0.87. 0.92, 0.92, 0.98, 1.32, 2.01(each 3H s, H3 of C-24, 25, 26, 27, 28, 23, 29, 30); 0.65 (m, 5a), 0.78-0.92 (m, 7a, 9a, 1b), 1.17 (15a), 1.22 (t-like, 11b), 1.32 (+, 6b), 1.32 (m, 19a), 1.32 (m, 13b), 1.44-146 (m, 7b, 20a, 15b, 6a) , 1.52 (1a), 1.55 (m, 2b), 1.72-1.73 (m, 2a, 17b, 20b, 11a), 1.95( dd, 7.4, 12.4, H-19b), 2.94 (dd like, 3a), 3.69 (m, 2a, 6a), Sugar moiety: 4.41 (d, J=6.0 Hz, H’-1), 2.93 (m, H0-2), 3.90( m, H0-3), 3.69 (m, H0-4), 2.94 (m, H0 a-5), 3.72( m, H0 b-5). ^13^C NMR (C_5_D_5_N; 150 MHz). 38.3(C-1), 26.9(C-2), 87.3(C-3), 37.5(C-4), 54.4(C-5), 17.8(C-6), 32.3(C-7), 45.7(C-8), 47.8(C-9), 35.6(C-10), 31.7(C-11), 67.2(C-12), 54.4(C-13), 40.4(C-14), 44.4(15), 64.2(16), 62.3(17), 45.7(C-18), 44.4(C-19), 36.3(C-20), 52.3(C-21), 213.6(C-22), 28.7(C-23),15.5(C-24), 14.7(C-25), 15.7(C-26), 19.7(C-27), 16.5(C-28), 19.7(C-29), 26.9(C-30), 105.6 (C-1’) , 72.6(C-2’), 83.1(C-3’), 68.7(C-4’), 65.1 (C-5’)

**Supplementary Figure 1:**

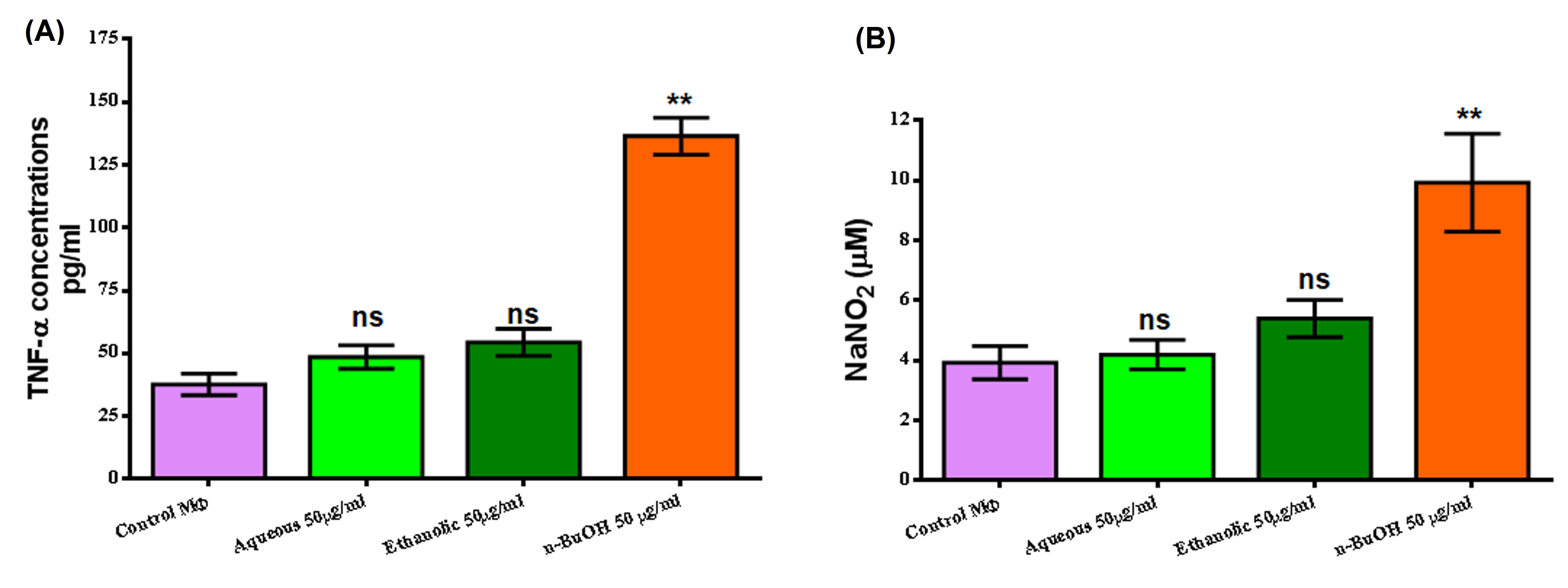

**Supple. Fig. 1. Evaluation of immunostimulation by fractions of *G. oppositifolius* MeOH extract.** The fractions, aqueous, ethanolic and n-BuOH (50μg/ml) were checked for altered TNF-α production after 24h treatment of RAW 264.7 MФs from the culture supernatant. (B) Extracellular NO production was monitored in the culture supernatant of treated (three fractions) and control after 24h by Griess reagent. All values are expressed as mean ± SEM from triplicate assays from three independent experiments (P values ≤ 0.05 (*) or ≤ 0.01 (**) vs. control).

**Supplementary Figure 2:**

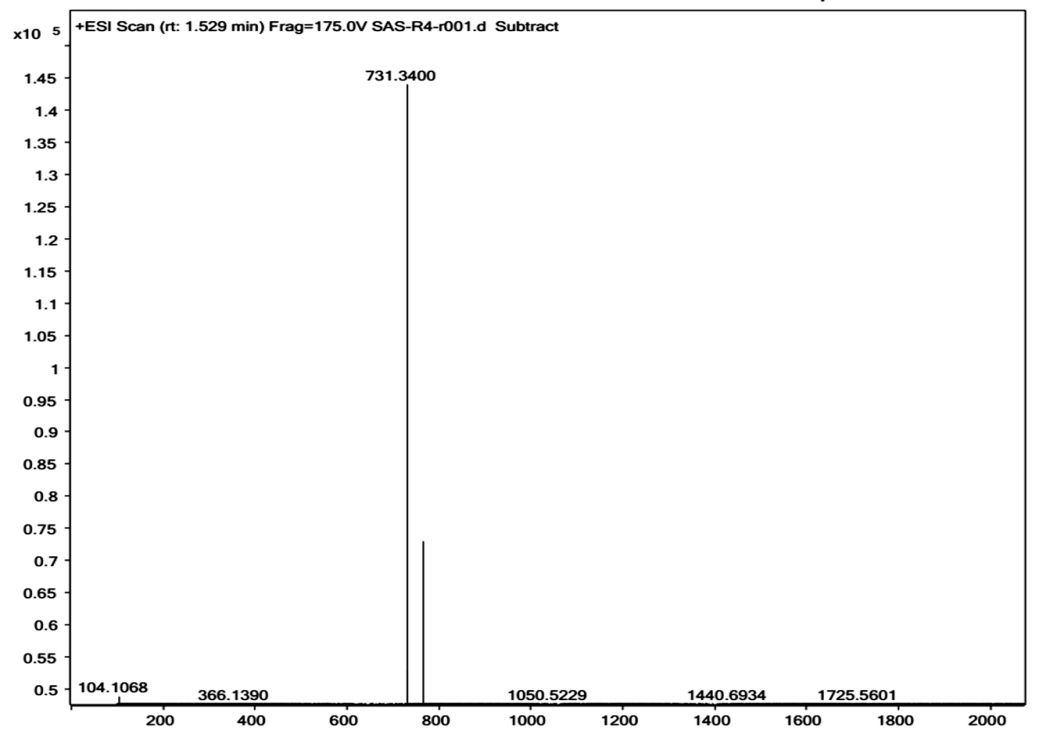

**Supple. Fig. 2. ESI-Mass Spectra of Compound 1 (Spergulin A)**

**Supplementary Figure 3:**

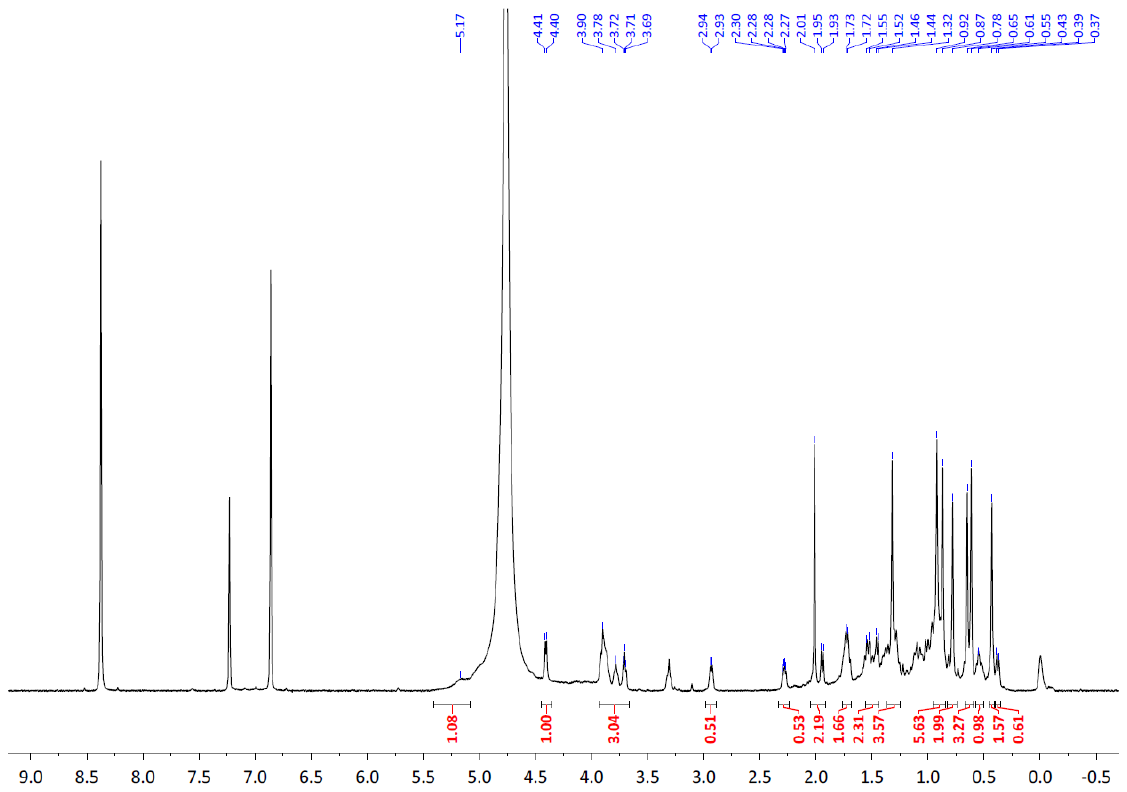

**Supple. Fig. 3. ^1^H NMR Spectra of Compound 1 (Spergulin A) in pyridine-d5**

**Supplementary Figure 4:**

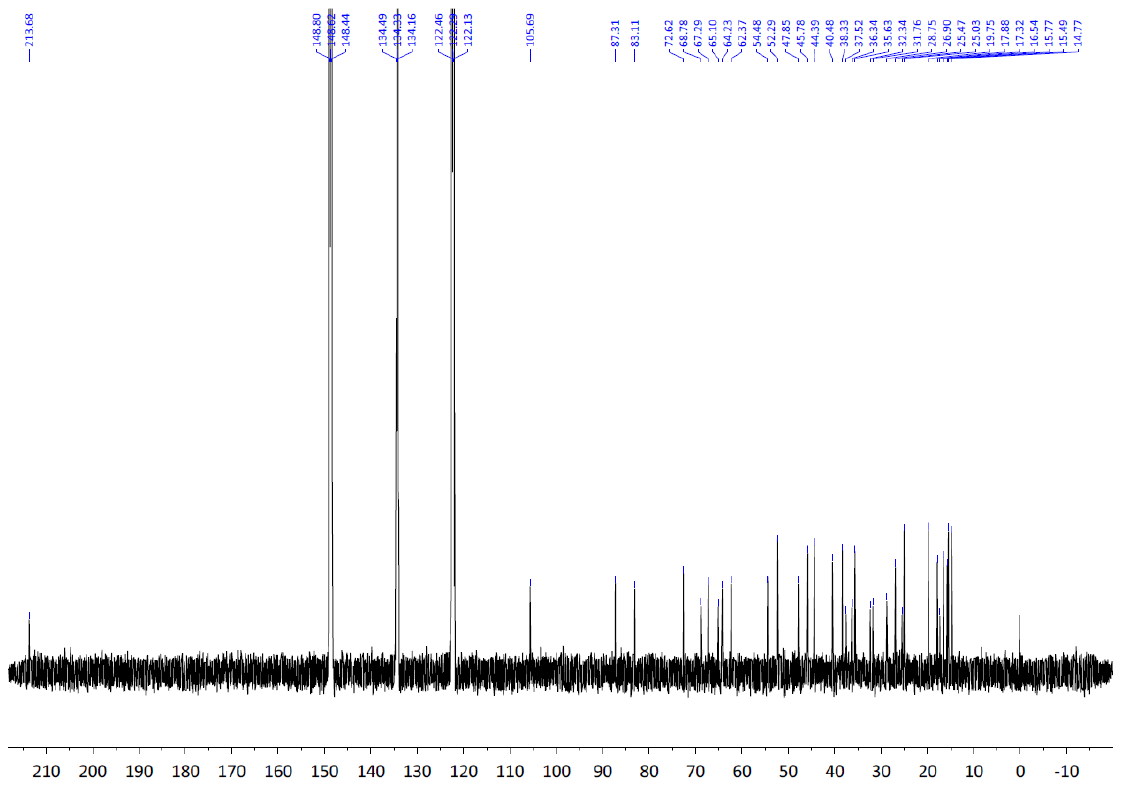

**Supple. Fig. 4. ^13^C NMR Spectra of Compound 1 (Spergulin A) in pyridine-d5**

**Supplementary Figure 5:**

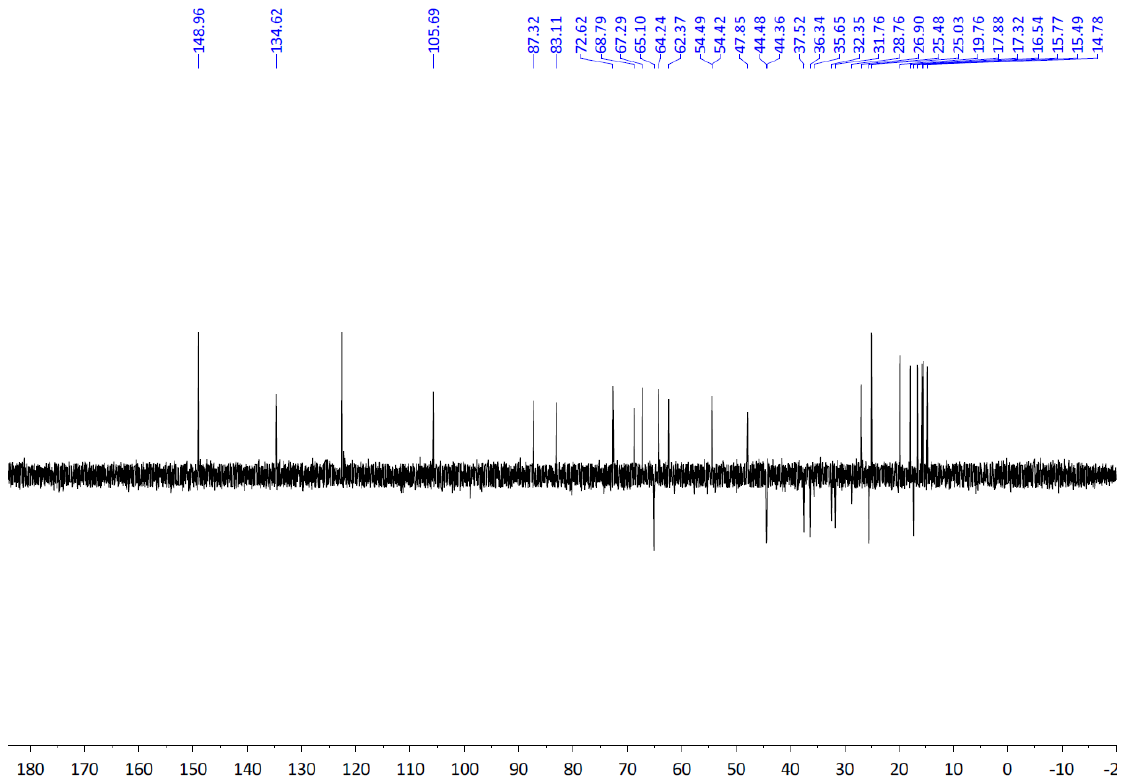

**Supple. Fig. 5. DEPT-135 of Compound 1 (Spergulin A) in pyridine-d5**

**(This pulse sequence produces a carbon spectrum with methyl (CH3) and methyne (CH) carbons are up. Methene (CH2) carbons are down)**

**Supplementary Figure 6:**

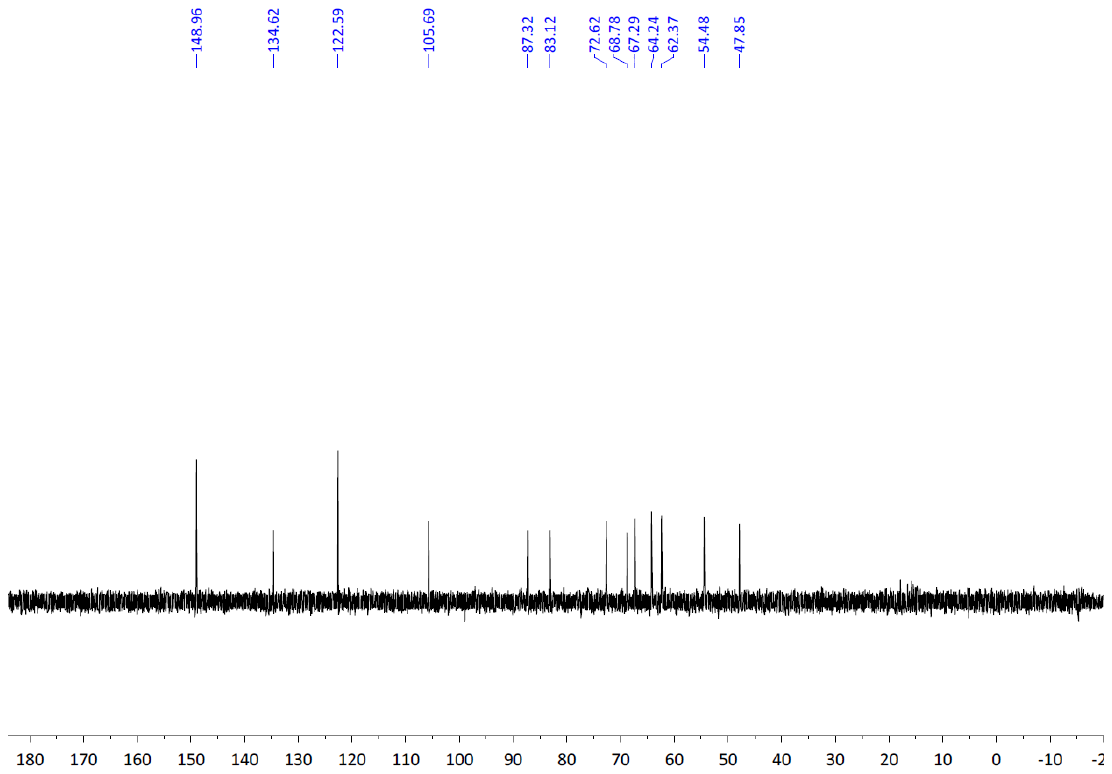

**Supple. Fig. 6. DEPT-90 of Compound 1 (Spergulin A) in pyridine-d5**

**(This pulse sequence produces a carbon spectrum containing only carbons with a single attached proton. Methyne, CH)**
